# Near-infrared spectroscopy calibration strategies to predict multiple nutritional parameters of pasture species from different functional groups

**DOI:** 10.1101/2021.07.31.454175

**Authors:** Karen L. M. Catunda, Amber C. Churchill, Sally A. Power, Ben D. Moore

## Abstract

Near-infrared reflectance spectroscopy (NIRS) has been used by the agricultural industry as a high-precision technique to quantify nutritional chemistry in plants both rapidly and inexpensively. The aim of this study was to evaluate the performance of NIRS calibrations in predicting the nutritional composition of ten pasture species that underpin livestock industries in many countries. These species comprised a range of functional diversity (C_3_ legumes; C_3_/C_4_ grasses; annuals/perennials) and origins (tropical/temperate; introduced/native) that grew under varied environmental conditions (control and experimentally induced warming and drought) over a period of more than 2 years (*n* = 2,622). A maximal calibration set including 391 samples was used to develop and evaluate calibrations for all ten pasture species (global calibrations), as well as for subsets comprised of the plant functional groups. We found that the global calibrations were appropriate to predict the six key nutritional quality parameters studied for our pasture species, with the highest accuracy found for ash (ASH), crude protein (CP), neutral detergent fibre and acid detergent fibre (ADF), and the lowest for ether extract (EE) and acid detergent lignin parameters. The plant functional group calibrations for C_3_ grasses performed better than the global calibrations for ASH, CP, ADF and EE parameters, whereas for C_3_ legumes and C_4_ grasses the functional group calibrations performed less well than the global calibrations for all nutritional parameters of these groups. Additionally, our calibrations were able to capture the range of variation in forage quality caused by future climate scenarios of warming and severe drought.

## 1. INTRODUCTION

Forage quality depends on plant nutritional composition, which influences digestibility and forage intake, and which are typically used as indicators of potential animal production (Ball et al., 2001; Coleman and Moore, 2003; Dumont et al., 2015). In view of this, evaluation of nutrient composition is essential for determining whether forage quality is adequate for animal production and in guiding livestock managers/farmers in determining how much forage and supplementation is needed to optimize food use efficiency for a particular animal and production goal (Ball et al., 2001). Traditionally, forage nutritional quality was evaluated by animal trials (e.g. *in vivo* and *in situ* digestibility with fistulated animals), however, this method has a number of constraints including high costs, labour, time investment and amount of the feed required in the trials. Consequently, this method is not suitable for examining large numbers or types of forage samples. Alternatives that have been used successfully to evaluate forage quality include proximate analysis of forage nutritional composition (nutrients and antinutrients), *in vitro* digestibility assays and near-infrared reflectance spectroscopy (NIRS) (Coleman and Moore, 2003).

Near-infrared reflectance spectroscopy has been embraced by the agricultural industry as a high-precision technique for measuring nutritional chemistry that is rapid, time-efficient, inexpensive, produces no chemical waste, and requires a small sample size and minimal preparation of samples (Abrams et al., 1987; Foley et al., 1998; Murray, 1993; Roberts et al., 2004). This is a non-destructive technique that uses the absorption and reflectance of nearinfrared light, and often visible wavelengths as well, from a sample to predict the chemical composition and other traits (e.g. digestibility, palatability etc.; Foley et al., 1998; Moore et al., 2010; Reddersen et al., 2013; Stuth et al., 2003). The reflectance spectrum of near-infrared light is influenced primarily by the nature of chemical bonds between hydrogen and carbon, hydrogen and nitrogen, and hydrogen and oxygen in each sample, and consequently by the nature and quantity of complex carbon and nitrogen-containing compounds, such as crude protein, fibre, and other plant constituents (Foley et al., 1998; Parrini et al., 2018; Smith et al., 2019). Based on these, concentrations can often be accurately predicted from the near-infrared reflectance spectra by developing standardized calibrations with samples of known nutritional composition (Foley et al., 1998). The process involves well-established statistical procedures used to develop, assess and improve predictive calibration equations for reflectance spectra based on reference values obtained by a variety of standard wet chemistry or other analytical techniques (Stuth et al., 2003). Importantly, accurate NIRS predictions of unknown samples depend on a calibration set (i.e. a large database) that is representative of the chemical and spectral variation encountered in the target population (Foley et al., 1998; Puigdomènech et al., 1997; Reddersen et al., 2013; Shenk and Westerhaus, 1991). After the development of reliable calibrations, NIR spectra of new samples can then be acquired and used for immediate quantification of multiple parameters, bypassing the need for wet chemistry analyses, and the associated expense of the latter.

Near-infrared reflectance spectroscopy has been shown to accurately predict forage nutritional quality, such as in studies involving mixed bulk samples of central European grasslands (Berauer et al., 2020), fresh samples from natural pastures in Italy (Parrini et al., 2019), warm-season legumes in the United States (Baath et al., 2020), and native and temporary grasses in the United Kingdom (Bell et al., 2018). However, few studies report on the establishment of NIRS calibration models for predicting multiple nutritional constituents, such as ash, crude protein, ether extract, neutral detergent fibre, acid detergent fibre and acid detergent lignin, and fewer still include calibrations using large sample sets comprised of multiple pasture species (Parrini et al., 2018; Parrini et al., 2019; cf. Norman et al., 2015; Norman et al., 2020). In addition, many calibration models for predicting nutritional composition do not capture variation across multiple species for a range of functional diversity and origins of pasture species, particularly growing under future environmental conditions such as climate change scenarios (Berauer et al., 2020).

In southern Australia, pasture systems are based on a diverse range of grasses and legumes, for which information on nutritional composition under a wide range of environmental (including climatic) conditions is limited (Howden et al., 2008; Lee et al., 2013; Norman et al., 2020; Norman et al., 2021). To address this knowledge gap, we used the Pastures and Climate Extremes (PACE) experimental facility to evaluate the nutritional responses of a wide range of pasture/rangeland species (including tropical/temperate, introduced/native, grasses/legumes) to year-round warming (including intensification of heatwaves) and extreme winter/spring droughts events. We specifically aimed to evaluate the performance of NIRS calibrations in predicting species’ nutritional composition under differing climatic conditions. The species used in this study comprised a range of functional diversity (C_3_ legumes; C_3_/C_4_ grasses; annuals and perennials) and origins (tropical and temperate; introduced and native) with a wide variation in concentrations of chemical constituents. Additionally, we tested different calibration strategies using combined datasets from all studied pasture species (global calibrations) and different independent datasets of plant species groups (plant functional group calibrations), in order to contribute to the improvement of NIRS calibrations and wider use of this technology in the research and development of livestock nutrition.

## 2. MATERIALS AND METHODS

### 2.1. Pasture sample collection

Representative samples of the pasture species were collected at the Pastures and Climate Extremes (PACE) experimental field facility and from an associated glasshouse study at the Hawkesbury Campus of Western Sydney University, at Richmond, NSW, Australia (S33.610, E150.740, elevation 25 m). The field site has a mean annual precipitation of 800 mm and mean annual temperature of 17.2°C, with monthly means peaking in January (22.9 C) and at their lowest in July (10.2°C) (Australian Government Bureau of Meteorology, Richmond-UWS Hawkesbury Station). At the field facility, the soil was a loamy sand with a volumetric water-holding capacity of 15 - 20%, pH of 5.7, plant available N of 46 mg/kg, plant available (Bray) P of 26 mg/kg and 1% soil organic carbon (more details are reported in Churchill et al., 2020). In a companion study using a glasshouse facility, two of the species from the field site (*Festuca arundinacea* and *Medicago sativa*) were grown using field soil. A detailed overview of the experimental facilities descriptions and pasture management are reported in Churchill et al. (2020; field), Catunda et al. (2021; glasshouse), and Zhang et al. (2021; glasshouse). The ten different pasture species used in this study include a range of plant functional groups (C_3_ legumes; C_3_/C_4_ grasses; annuals and perennials) and origins (tropical and temperate; introduced and native) and are commonly used as forage in grasslands in south-eastern Australia and, with the exception of the grass *Rytidosperma caespitosum*, internationally (**Table 1**). Plants were grown under a wide range of environmental conditions, including warming and/or drought treatments; (Churchill et al., 2020; Catunda et al., 2021), while other conditions were held constant (soils, fertilization, pests, etc.). These experimentally manipulated conditions maximised the range of variation in the concentrations of chemical components (Catunda et al., 2021).

**Table 1.** Information about pasture species, climate conditions and sample number included in the study.

| Type | Photosynthetic<br>Pathway | Common name | Latin name (cultivar) | Origin | Lifecycle | Climate conditions<sup>#</sup> | Sample<br>number |
| --- | --- | --- | --- | --- | --- | --- | --- |
| Legumes* | C<sub>3</sub> | Biserrula | *Biserrula pelecinus* (Casbah) | Temperate, introduced | Annual | Drought | 139 |
|  | C<sub>3</sub> | Lucerne | *Medicago sativa* (SARDI 7 series 2) | Temperate, introduced | Perennial | Drought and warming | 542 |
|  | C<sub>3</sub> | Sub-clover | *Trifolium subterraneum* (Campeda) | Temperate, introduced | Annual | Drought and warming | 29 |
| Grasses | C<sub>3</sub> | Phalaris | *Phalaris aquatic* (Holdfast GT) | Temperate, introduced | Perennial | Drought and warming | 332 |
|  | C<sub>3</sub> | Ryegrass | *Lolium perenne* (Kidman) | Temperate, introduced | Annual | Drought | 92 |
|  | C<sub>3</sub> | Tall Fescue | *Festuca arundinacea* (Quantum II MaxP) | Temperate, introduced | Perennial | Drought and warming | 429 |
|  | C<sub>3</sub> | Wallaby | *Rytidosperma caespitosum* | Temperate, native | Perennial | Drought and warming | 185 |
|  | C<sub>4</sub> | Digit | *Digitaria eriantha* (Premier) | Tropical, introduced | Perennial | Drought | 300 |
|  | C<sub>4</sub> | Kangaroo grass | *Themeda triandra* | Tropical, native | Perennial | Drought and warming | 417 |
|  | C<sub>4</sub> | Rhodes | *Chloris gayana* (Katambora) | Tropical, introduced | Perennial | Drought | 157 |

\*Legumes received appropriate rhizobium inoculant during sward establishment: ALOSCA granular inoculant for *Biserrula* (Group BS; ALOSCA Technologies, Western Australia, Australia); Easy Rhiz soluble legume inoculant and protecting agent to *Lucerne* (Group AL; New Edge Microbials, New South Wales, Australia); and NoduleN for the *Sub-clover* (Group C; New Edge Microbials).
<sup>#</sup>A detailed overview of the climate conditions is reported in Churchill et al. (2021; field), Catunda et al. (2021; glasshouse) and Zhang et al. (2021; glasshouse).

Plant samples from the field were collected throughout a period of 2 years and 4 months (from November 2017 to March 2020) in regular harvests of aboveground biomass based on cut and carry recommendations used by local farmers (Clements et al., 2003); perennial species were harvested 3-5 times per year and annual species 2-3 times (Clark et al., 2016). Samples from the glasshouse study were collected in June and August 2018 following the same harvest protocol adopted in the field. Swards were cut at 5 cm above the soil surface and weighed (fresh weight). The representative samples were composed of leaves, stems/tillers, and flowers when present, as well as a mixture of both live and dead material as present. Weeds were removed prior to the collection of near-infrared reflectance spectra and wet chemical analysis of a subset of the target species’ material. A total of 2,622 samples (2,238 from the field and 384 from the glasshouse) were collected and used for evaluation as described below (**Table 1**).

### 2.2. Sample preparation and spectral data collection

Forage samples were either immediately frozen (−18°C) and later freeze-dried (for the glasshouse facility) or microwaved at 600 W for 90 seconds to deactivate enzymes (Landhäusser et al., 2018), followed by oven-drying at 65°C for 48 h (for the field facility). To homogenize samples, dried plants were ground through a 1-mm screen in a laboratory mill (Foss Cyclotec Mill, Denmark) and stored in airtight plastic containers in the dark at room temperature prior to the collection of near-infrared reflectance spectra and wet chemical analysis. For the nitrogen analysis, plant samples were ground further, using a ball-mill to produce a fine powder (Retsch® MM200; Hann, Germany).

Near-infrared spectra were collected on a FOSS XDS Rapid Content™ Analyzer with XDS near-infrared technology (FOSS Analytical, Hilleroed, Denmark). All samples were analysed in the range of 400-2500 nm with a spectral resolution of 0.5 nm. Spectra were acquired with ISIscan™ Routine Analysis Software (Foss, Denmark). Samples were repacked and scanned in duplicate, and the average spectrum for each sample was used for subsequent calibrations and predictions. A subset of 391 representative samples was selected for determining nutritional composition by wet chemistry using the ‘select’ function in the software WinISI 4.8.0 (FOSS Analytical A/S, Denmark). The selected samples for wet chemical analyses were representative across sample collection periods, field and glasshouse settings, functional diversity and species’ origin. Furthermore, the subset covered the range of spectral variation in the full scanned population of plant samples, summarized by a principal component analysis to minimize spectral redundancy.

### 2.3. Wet chemical analysis

Samples were subjected to analyses of dry matter (DM) and ash (ASH) according to the official methods and procedures for animal feed outlined by the Association of Official Analytical Chemists (AOAC, 1990). Nitrogen (N) concentration was determined from ∼ 100 mg of sample using an automated combustion method with a Leco TruMac CN analyzer (Leco Corporation, USA). Crude protein (CP) concentration was then calculated by applying a 6.25 conversion factor to the N concentration (AOAC, 1990). Ether extract (EE) was determined according to the American Oil Chemists’ Society (AOCS) high-temperature method using petroleum ether (B.P. 40-70°C) and the Soxhlet method (Buchi 810 Soxhlet Multihead Extract Rack, UK). Fibre fractions were determined with an ANKOM Fibre Analyzer (model 200, ANKOM® Technology, NY, USA) with use of neutral and acid detergent solutions and correction for dry matter content (Goering and Van Soest, 1970). The samples were analysed for neutral detergent fibre (NDF), acid detergent fibre (ADF) and acid detergent lignin (ADL) by the sequential method of Van Soest and Robertson (1980). Sodium sulphite and α-amylase were added to the solution for NDF determination. Each sample was analysed in duplicate and nutrient concentrations were expressed on a DM basis (as a percentage).

### 2.4. Calibration development and statistics

NIRS calibration models were established using WinISI software version 4.8.0 (FOSS Analytical A/S, Denmark). We adopted two calibration strategies (**Figure 1)**: 1) global and 2) plant functional group-based. For the global calibration models development, samples from all pasture species were subjected to wet chemistry analysis (n = 391), and randomly assigned to either a calibration set (n = 313 samples) or an external validation set (n = 78 samples). We also investigated the performance of dedicated calibration models for each of three functional group categories: (1) C_3_ legumes, (2) C_3_ grasses and (3) C_4_ grasses (**Figure 1**). For each of these plant functional group models, 80% of the chemistry/spectra pairs were used for the calibration set and the remainder were kept for external validation.

**Figure 1.**
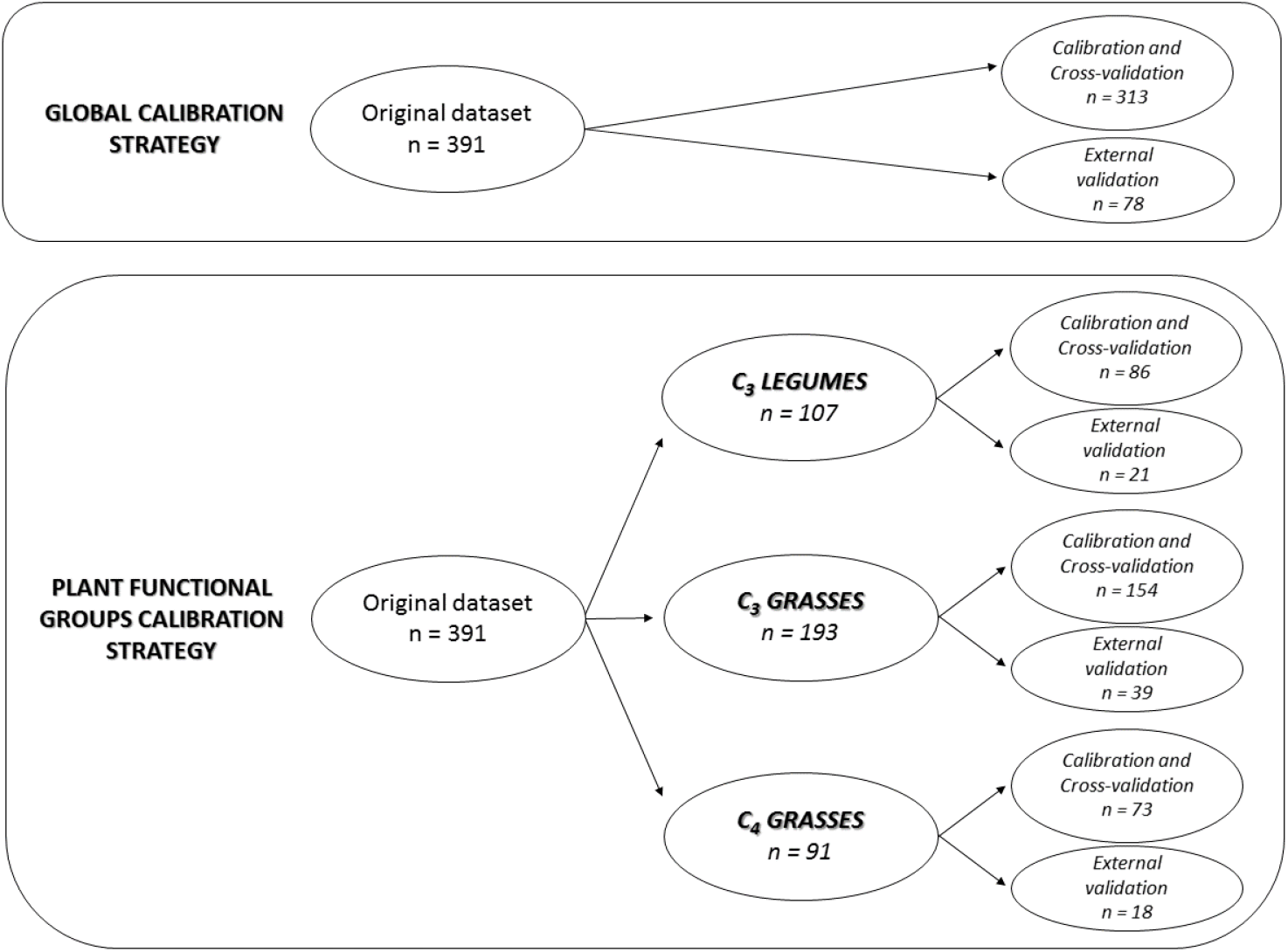
Summary of the datasets, calibration and validation processes used in two calibration strategies: a global calibration that included all species and dedicated plant functional group calibrations that modelled C_3_ legumes, C_3_ grasses and C_4_ grasses separately.

Predictive equations based on samples analysed with wet chemistry were then developed using modified partial least-squares regression (MPLS), with cross-validation to prevent overfitting of models (Shenk and Westerhaus, 1991). The best performing equations were selected by testing a range of scattering pre-treatment options with different derivative gaps and smoothing using the software WinISI 4.8.0. The variety of scattering pre-treatments tested included none, standard normal variate + detrend, standard normal variate only, detrend only, standard multiplicative scatter correction, and weight multiplicative scatters correction in order to reduce the influence of the sample particle size on NIRS spectra and the path-length variations. The different derivative mathematical pre-treatments tested for each calibration equation to decrease noise effects were coded as follows: “1,4,4,1,” “1,6,4,1,” “1,8,6,1,” “2,4,4,1,” “2,6,4,1,” “2,8,6,1,” “2,10,10,1,” and “3,10,10,1,” where the first digit is the number of the derivative, the second one is the gap over which the derivative is calculated, the third one is the smoothing segment and the last one is the secondary smoothing segment. For the six key nutritional parameters, the best models were selected on the basis of the highest coefficient of determination of calibration (R^2^) and the internal cross-validation (one minus the variance ratio, 1-VR), along with the lowest standard error of calibration (SEC) and internal cross-validation (SECV), and the smallest difference between SEC and SECV; this was achieved using the software WinISI 4.8.0 (Andueza et al., 2011; Norman et al., 2020). To compare the predictive ability of calibration equations among different parameters, the correlation coefficient of actual and predicted values (R^2^) obtained in the external cross-validation set was evaluated. In addition, to support data interpretation and allow comparison with other studies, we calculated the ratio performance deviation (RPD) by dividing the standard deviation of the reference data by the standard error of prediction (Williams and Sobering, 1996; Williams, 2014). All scatter plots were created using R software, version 4.0.0 (R Core Team, 2020).

## 3. RESULTS AND DISCUSSION

The ranges and means for the six nutritional parameters (ASH, CP, EE, NDF, ADF and ADL) of pasture species used in the calibration models are presented in **Table 2**. Our wet chemistry results reveal a broader range of values than those reported in previous studies with forage species (Andueza et al., 2011, Parrini et al., 2019, Parrini et al., 2018), allowing our NIRS calibration models in term to be applied to samples spanning a broader range of quality values. This may reflect the fact that as well as comprising a broad range of functional diversity and species origins, our plants were also grown under very varied environmental conditions, including control conditions as well as warming and severe droughts, imposed individually and in combination. Warming treatments included continuous + 3°C above-canopy temperature (field experiment; Churchill et al., 2020) and + 4°C above ambient (glasshouse experiment; Catunda et al., 2021; Zhang et al., 2021). Drought treatments consisted of winter and spring periods of 60% rainfall reduction (field experiment), and 60% soil water holding capacity reduction (glasshouse experiment). These climate treatments represented the extreme climate change predictions for average surface air temperatures and winter/spring droughts for the study region (CSIRO and BoM, 2015; CSIRO, 2020).

**Table 2.** Details of treatment of spectra, calibration and validation sets, and performance statistics of the global and C_3_ grasses calibration models for nutritional parameters (expressed as percentage of dry matter) of pasture species.

| Calibrations | Parameter | Range (%) | Mean (%) | Wavelengths (nm) | Mathematical treatment* | Calibration |  |  |  |  |  | Validation |  |  |  |
| --- | --- | --- | --- | --- | --- | --- | --- | --- | --- | --- | --- | --- | --- | --- | --- |
|  |  |  |  |  |  | N | SEC | R<sup>2</sup>c | SECV | 1-VR | RPD | N | SEP | R<sup>2</sup>v | RPD |
| Global | ASH | 3.1 - 19.0 | 8.1 | 400-2492 | 2,8,6,1 | 307 | 0.41 | 0.96 | 0.73 | 0.88 | 5.0 | 78 | 0.91 | 0.89 | 3.0 |
|  | CP | 1.7 - 32.0 | 12.0 | 700-2492 | 3,10,10,1 | 309 | 0.89 | 0.97 | 1.13 | 0.95 | 5.9 | 78 | 1.10 | 0.95 | 4.5 |
|  | EE | 0.1 - 7.6 | 2.9 | 400-2492 | 2,6,4,1 | 306 | 0.54 | 0.92 | 0.89 | 0.79 | 3.6 | 78 | 0.99 | 0.72 | 2.0 |
|  | NDF | 22.0 - 84.2 | 58.2 | 400-2492 | 2,8,6,1 | 311 | 1.78 | 0.98 | 2.50 | 0.96 | 7.1 | 78 | 2.34 | 0.97 | 5.9 |
|  | ADF | 15.8 - 54.7 | 31.9 | 400-2492 | 2,4,4,1 | 308 | 1.41 | 0.94 | 1.80 | 0.89 | 4.2 | 78 | 2.06 | 0.91 | 3.3 |
|  | ADL | 0.3 - 18.3 | 6.1 | 400-2492 | 3,10,10,1 | 309 | 1.14 | 0.85 | 1.51 | 0.73 | 2.6 | 78 | 1.55 | 0.72 | 2.0 |
| C<sub>3</sub> grasses | ASH | 3.4 - 18.9 | 8.7 | 400-2492 | 3,10,10,1 | 151 | 0.35 | 0.98 | 0.87 | 0.87 | 7.1 | 39 | 0.73 | 0.93 | 3.8 |
|  | CP | 2.8 - 27.8 | 11.5 | 700-2492 | 2,4,4,1 | 153 | 0.30 | 0.99 | 0.73 | 0.97 | 10.0 | 39 | 0.87 | 0.97 | 5.9 |
|  | EE | 0.1 - 7.6 | 3.3 | 400-2492 | 2,4,4,1 | 150 | 0.43 | 0.95 | 0.80 | 0.83 | 4.5 | 39 | 0.98 | 0.75 | 2.0 |
|  | ADF | 19.1 - 46.1 | 30.6 | 400-2492 | 2,4,4,1 | 153 | 0.83 | 0.98 | 1.67 | 0.92 | 7.1 | 39 | 1.38 | 0.95 | 4.5 |

*ASH: ash; CP: crude protein; EE: ether extract; NDF: neutral detergent fibre; ADF: acid detergent fibre; and ADL: acid detergent lignin.*
*\*Mathematical treatment describes the approach used for spectral analysis (stored as log(1/reflectance)). The first two numbers describe the* *derivative used, the third and fourth numbers indicate the degrees of primary and secondary smoothing performed on the derivative. Thus 2,4,4,1* *indicates that the second derivative was calculated with a gap size of 4 nm and that a maximal primary smooth but no secondary smooth was used.*
*N: The number of samples in the calibration or validation set; SEC: standard error of calibration; R<sup>2</sup>c: coefficient of determination of calibration;* *SECV: standard error of the internal cross validation; 1-VR: coefficient of determination of the internal cross-validation; RPD: ratio of standard* *error of performance: standard deviation of calibration or validation; SEP: standard error of prediction; and R<sup>2</sup>v: coefficient of determination of* *validation.*

Overall, our calibrations accurately predicted the nutritional composition of pasture biomass across a range of species and plant functional groups, which accords with previous studies predicting forage quality by NIRS (Andueza et al., 2011; Smith et al., 2019). Optimal calibrations were obtained using different calibration parameters for each of the key nutritional parameters, including different mathematical spectral pre-treatments for each constituent, although standard normal variate + detrend was the most effective scattering pre-treatment in all cases, as also reported by Barnes et al. (1989).

The samples in our study, across all pasture species, were first value-predicted for each nutritional parameter using equations developed by the global calibrations (**Figure 1**), then samples from each plant species group (C_3_ legumes, C_3_ grasses, and C_4_ grasses) were separated and predicted using their respective plant functional group calibrations in order to improve the results that were obtained with the global calibrations. From this step, C_3_ legumes and C_4_ grasses functional group calibrations did not improve the accuracy of any nutritional parameters compared to results obtained using the global calibrations, however, the C_3_ grasses functional group calibration improved predictions, relative to the global calibrations, for some nutritional parameters (ASH, CP, EE and ADF). Norman et al. (2015; 2020) investigated the use of NIRS calibrations to predict the nutritional value of grasses, legumes and forbs, concluding that separating taxonomically similar species into groups, did not lead to more accurate predictions than broad, mixed species calibrations. The best-performing calibration models obtained in our study for each nutritional parameter based on optimal wavelengths, mathematical pre-treatments, scattering processing and regression method are summarized in **Table 2** (global and C_3_ grasses calibration models).

Most models used wavelengths from 400-2500 nm, with the exception of CP which used wavelengths from 700-2500 nm (**Table 2**). Among the chemical constituents, different chemical bonds absorb at different wavelengths, thus identifying these regions located in the spectrum contributes to better estimation of the concentration of these nutritional parameters (Foley et al., 1998). In addition, the best calibration statistics in our study were found using the second derivative mathematical treatment for most of the parameters calibration models, whereas the third derivative treatment was best for CP and ADL in the global calibration and ASH for the C_3_ grasses calibration models (**Table 2**).

The global calibrations had high accuracy and predictive power in both internal (cross) and external validation for ASH, CP, NDF and ADF (**Table 2; Figure 2**). The accuracy was lower for EE and ADL than other parameters examined in this study (**Table 2**; **Figure 2**), and this may be due to the occurrence of samples with low concentrations of these constituents (**Table 2**). The low concentrations found here could cause changes in the spectrum wavelength and absorption, making it difficult to measure using NIRS, compared to the other nutritional parameters, as reported by Roberts et al. (2004). Only a few studies have reported NIRS calibrations for EE and ADL nutritional parameters, making comparison and clarification challenging; this may reflect a general difficulty in achieving satisfactory calibrations. Berauer et al. (2020) in a study with 512 (calibration set) bulk samples of European species-rich montane pastures, reported similar predictions for EE (R^2^c = 0.86, R^2^v = 0.73). In contrast, Parrini et al. (2018) presented calibration models using 105 bulk pasture samples in their calibration set collected from Tuscany (Italy) that were able to predict EE and ADL with higher accuracy than in our study (for both parameters R^2^c =0.99, R^2^v =0.98). In the latter case, the small number of samples used may have contributed to low variability of values between samples used in the calibration sets and lower errors in the associated predictions.

**Figure 2.**
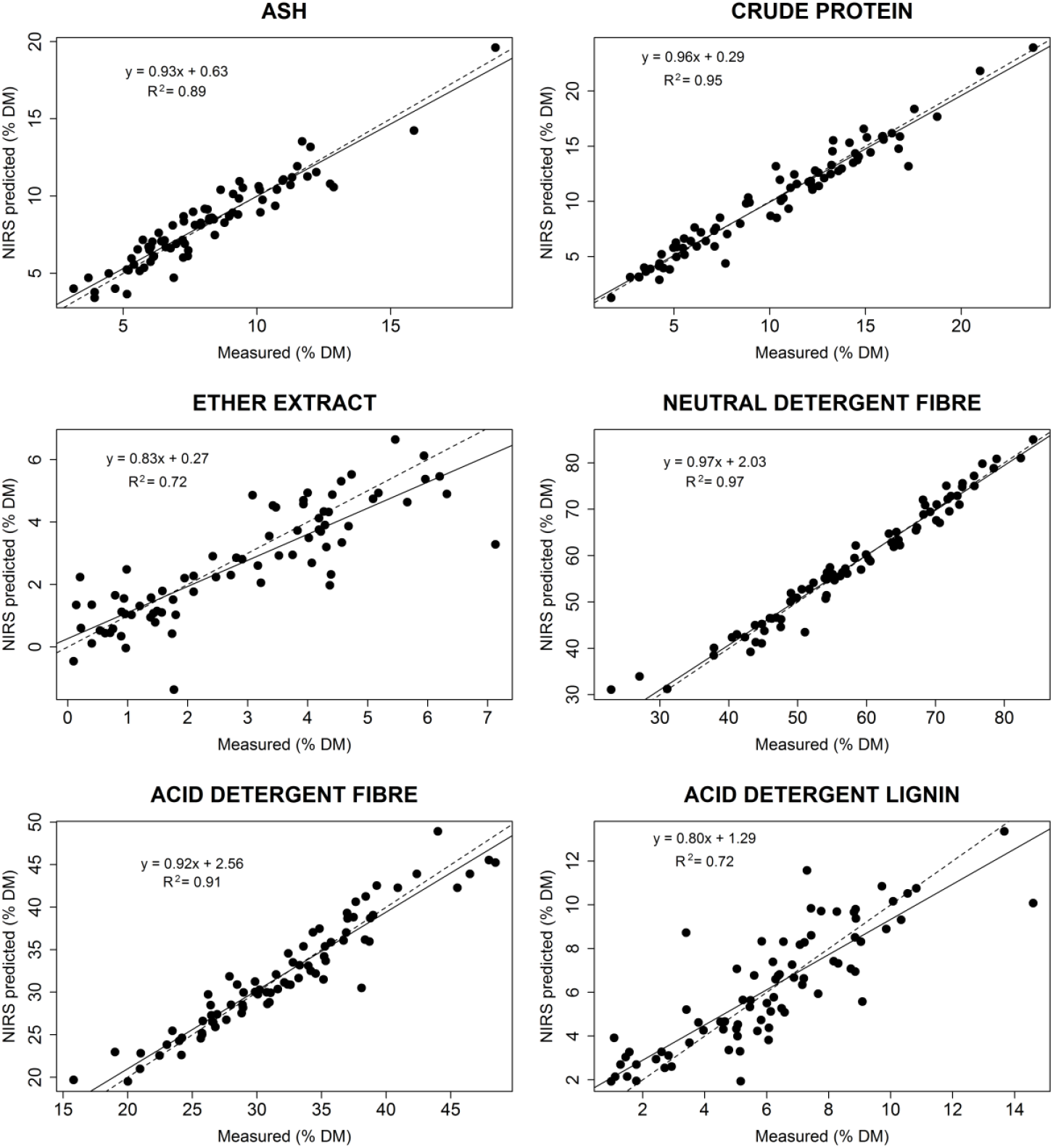
Relationships between values measured in laboratory analysis and values predicted by NIRS in the external validation set using the global calibrations for nutritional parameters (ash, crude protein, ether extract, neutral detergent fibre, acid detergent fibre and acid detergent lignin), expressed as percentage of dry matter (% DM). Solid lines indicate ordinary leastsquares linear regressions, and dashed lines show a 1:1 relationship.

Plant functional groups are widely used to describe trait variation within and across plant communities (Thomas et al., 2019). Despite the widespread use of plant functional groups to describe common plant morphological, physiological, biochemical and phenological traits (Pérez-Harguindeguy et al., 2013), in our study calibration models based on plant functional groups were only superior to the global calibrations for C_3_ grasses. The predictions for C_3_ grass samples were superior using the C_3_ grass functional group calibration models for ASH, CP, EE and ADF (**Table 2, Figure 3**).

**Figure 3.**
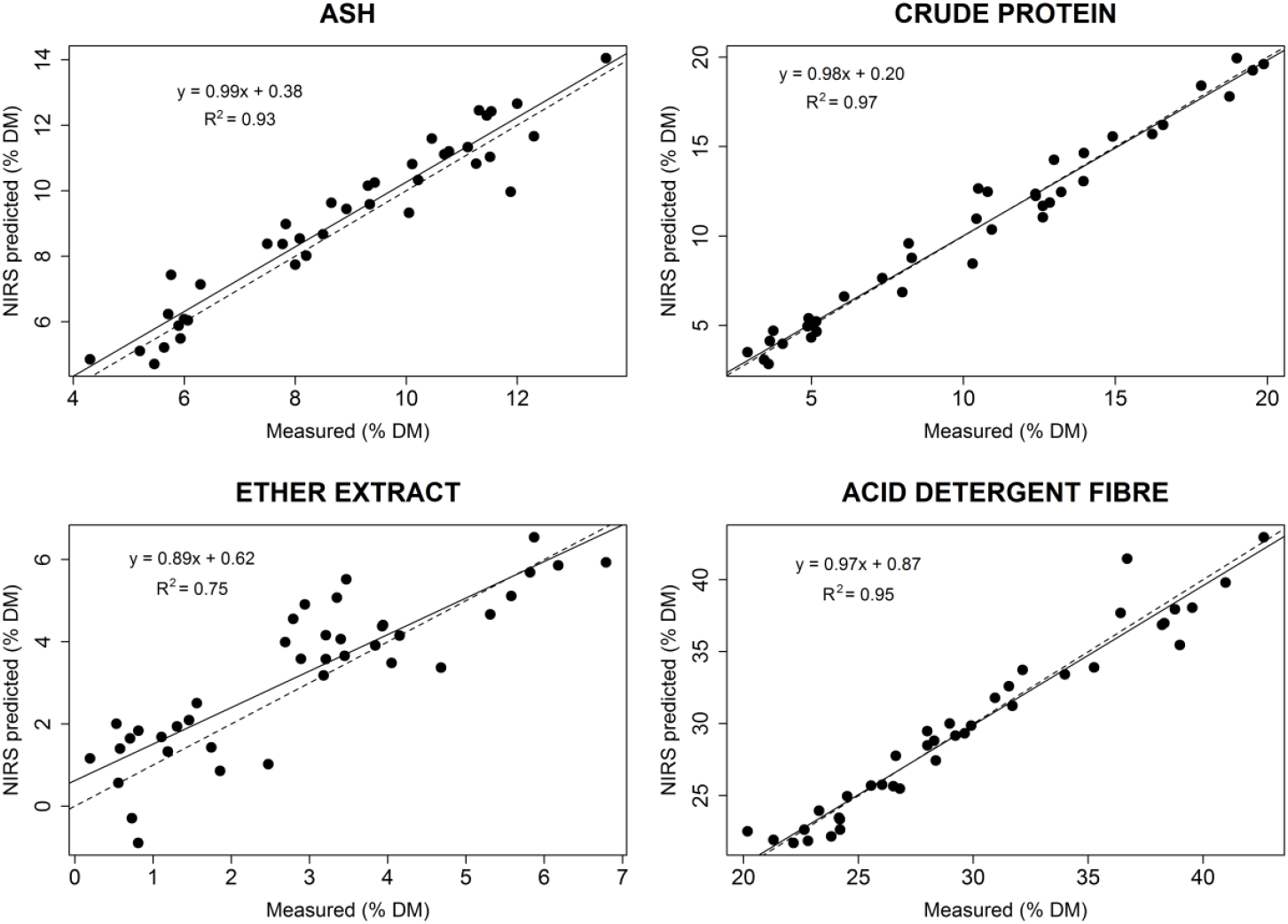
Relationships between values measured in laboratory analysis and values predicted by NIRS in the external validation set using the C_3_ grass functional group calibrations for nutritional parameters (ash, crude protein, ether extract, acid detergent fibre) of C_3_ grasses, expressed as percentage of dry matter (% DM). Solid lines indicate ordinary least-squares linear regressions and dashed lines show a 1:1 relationship.

In order to evaluate the accuracy of a calibration model and to allow standard comparison with other studies, we calculated the RPD (ratio performance deviation), a non-dimensional statistic for the quick evaluation and classification of NIR spectroscopy calibration models which has been widely used in NIRS studies (Williams and Sobering, 1996; Williams, 2014). In our study, RPD values from the validation set for ASH, CP, NDF and ADF (**Table 2**) were acceptable for quality control, ranging from 3 for ASH (global calibrations) to 6 for the NDF (global calibrations) and CP (C_3_ grasses calibrations) equations, with respectively “good” to “excellent” classification according to Williams (2014). The high accuracy found in our study for predictions of NDF and CP may be related to the wide range of concentrations in biomass samples included in this study, as also reported by Andrés et al. (2005) in predictions of forage species from Leon (Spain). RPD values (validation set) for EE (global and C_3_ grasses calibration models) and ADL (global calibrations) were lower (both RPD = 2) than the other parameters, although these are considered acceptable for screening purposes (Williams, 2014). For Australian forage species, Norman et al. (2020) developed calibration models that included 3 years of data collection of forage species, found RPD values for validation datasets of CP, NDF and ADF that are comparable to those we report here; however, they did not report EE and ADL parameters so comparison for these is not possible. Other studies with multiple forage species showed lower NIRS predictive ability than our models for specific nutritional parameters such as ASH (Andueza et al., 2011; Berauer et al., 2020; Norman et al., 2020), CP (Andueza et al., 2011; Berauer et al., 2020), EE (Berauer et al., 2020), NDF and ADF (Smith et al., 2019).

Our NIRS calibrations provided satisfactory accuracy (predictive power) to be able to detect changes in forage nutritional quality associated with differences in phenology and warming and/or drought scenarios over the 2 years of sample collections. These characteristics qualify our calibrations to assess the effects of seasonality on forage quality and suggest the feasibility of future development of real-time in-field NIRS monitoring approaches to monitor seasonal and interannual changes in nutrient concentrations of pasture species (Bell et al., 2018; Murphy et al., 2021). These abilities will help farmers/ industry to assist with regular feed management decision-making, including in the face of climate change and associated climate extremes, such as are increasingly being experienced across Australia and, indeed, worldwide.

## 4. CONCLUSION

This study showed that global NIRS calibrations for a diverse range of pasture species were able to predict multiple key nutritional parameters. Predictions of nutritional metrics for C_3_ grass biomass were improved by using a plant functional group-specific calibration models for ash, crude protein, ether extract and acid detergent fibre, whereas those for C_3_ legumes and C_4_ grasses were accurately predicted using the global calibrations. In addition, our calibrations explicitly capture the range of variation in forage quality brought about by warming and drought treatments in this suite of pasture species. High quality, accurate NIRS calibrations are an essential tool to help rapidly track/monitor forage quality changes in response to management interventions and climate conditions, consequently improving pasture management practice in the future.

## ACKNOWLEDGEMENTS

We are grateful to all the team members in Pasture and Climate Extreme (PACE) project specially Chioma Igwenagu, Gil Won Kim, Haiyang Zhang, Kathryn Fuller, Manjunatha Chandregowda and Vinod Jacob, as also WSU summer scholars: Alexandra Boyd, Minh Doan, Samantha Weller, Shania Therese Didier Serre and Ben Capel. This work was supported by funding from the Meat & Livestock Australia Donor Company (P.PSH. 0793), Dairy Australia (C100002357) and Western Sydney University. The authors would like to thank the technical team at Western Sydney University for technical support. The authors declare that there are no conflicts of interest regarding the publication of this article.

